# The cysteine-rich domain of SEP15, a selenoprotein co-chaperone of the ER chaperone, UDP-glucose:glycoprotein glucosyltransferase, adopts a novel fold

**DOI:** 10.1101/2025.08.11.669745

**Authors:** Robert V. Williams, Kevin P. Guay, Owen Hurlbut Lesk, Daniel N. Hebert, Lila M. Gierasch

**Affiliations:** Department of Biochemistry & Molecular Biology, University of Massachusetts Amherst, Amherst, MA 01003; Department of Chemistry, University of Massachusetts Amherst, Amherst, MA 01003; Program in Cell and Molecular Biology, University of Massachusetts Amherst, Amherst, MA 01003

**Author notes:** Deceased (Daniel N. Hebert).

**Keywords:** Endoplasmic reticulum, protein folding, glycoprotein, selenoprotein, disulfide

## Abstract

Proteins targeted to the secretory pathway are involved in a myriad of biological processes but can only do so when properly folded. Within the endoplasmic reticulum, glycoprotein folding is regulated by the enzyme UDP-glucose:glycoprotein glucosyltransferase (UGGT) and its oxidoreductase partner, the 15-kDa selenoprotein (SEP15 aka SELENOF). The interaction between these two chaperones is poorly understood, limiting understanding of their function. SEP15 is comprised of two domains, a C-terminal thioredoxin-like domain, the structure of which has been reported (PDB 2A4H), and an approximately 50-residue long N-terminal cysteine-rich domain (CRD), of unknown structure. Here, we use a combination of AlphaFold structural predictions and NMR spectroscopy to elucidate the structure of the SEP15 CRD, which mediates the interaction with UGGT. These data reveal that this domain forms a previously undescribed helical fold stabilized by three disulfide bonds between residues C10-C42, C21-C43, and C24-C39. Furthermore, our results validate our reported model of the UGGT/SEP15 complex and lay the foundation for future studies of its interaction with glycoprotein substrates.

## Introduction

Roughly one-third of the human proteome is targeted to the secretory pathway (1). These proteins are translated directly into the endoplasmic reticulum (ER) lumen, where they fold to their mature forms. Several chaperone systems assist in this process, one of which, unique to the ER, is the lectin chaperone network (2). Folding assistance by the lectin chaperones is directed, in part, by N-linked glycan modifications. These carbohydrates are added as a Glc_3_Man_9_GlcNAc oligosaccharide by *en bloc* transfer via oligosaccharyltransferase (OST) to Asn residues within N-X-S/T/C sequence motifs. The A form of OST (OST-A) acts co-translationally, and shortly after addition the first two, terminal Glc residues are cleaved by a-glucosidase I and II (GlsI, GlsII). The resulting monoglucosylated glycan is recognized by the lectin chaperones, calnexin (CNX) and calreticulin (CRT), which also associate with co-chaperones to promote proper folding.

Following release from CNX/CRT the remaining Glc residue is cleaved by a second action of GlsII. At this point the glycoprotein has reached a checkpoint. If folded, it may traffic to its final cellular destination. However, if not properly folded, it interacts with the enzyme UDP-Glc:glycoprotein glucosyltransferase (UGGT), which will regenerate the monoglucosylated glycan and allow for the glycoprotein to re-engage CNX/CRT for additional rounds of folding.

A less well-understood aspect of UGGT’s roles in the ER is its partnership with a small, selenoprotein co-chaperone: the 15-kDa selenoprotein (SEP15, aka SELENOF). All SEP15 is in a stable complex with UGGT, but not all UGGT has a bound SEP15. SEP15 contains a single selenocysteine (Sec or U) residue, the 21^st^ amino acid in a redox-active C-G-U motif within its C-terminal domain, which adopts a minimal thioredoxin fold (3). SEP15 is one of only 25 selenocysteine-containing proteins in the human proteome, and the special properties of the Sec residue are likely exploited in SEP15 function. Importantly, the N-terminal Cys-rich domain (CRD) of SEP15 mediates its interaction with UGGT (4). Up to the present, no experimental structures have been obtained for the CRD.

SEP15 plays a critical but poorly understood role in ER protein quality control. Expression of SEP15 is regulated by the unfolded protein response (5). It has been proposed that SEP15 targets a subset of UGGT clients rich in disulfide bonds (6). At the physiological level, *Sep15* knockout mice developed cataracts at an early age and displayed elevated oxidative stress in the liver (7). Better understanding of the molecular basis of the interaction between UGGT and SEP15 will enhance understanding of ER protein homeostasis and aid future efforts at designing therapeutics that modulate it.

Previously, we described an AlphaFold prediction of the UGGT1/SEP15 complex, and we validated the model by complex-disruptive mutations on the UGGT surface where SEP15 was proposed to bind (8). The model of the UGGT/SEP15 complex provided a first pass structure of SEP15. However, the SEP15 model that emerged from this study was characterized by a disulfide pairing pattern that differed from a previous report in the literature (9). Consequently, we have now carried out a direct structural determination of the SEP15 CRD using NMR spectroscopy. The results are consistent with an AlphaFold prediction and show that the CRD adopts a novel helical fold stabilized by three disulfide bonds in a pattern that differs from that in the previous report. Moreover, the experimental NMR-based CRD structure is consistent with that in the model of the UGGT/SEP15 complex.

## Results

### AlphaFold3 predicts a helical fold for SEP15 CRD

As observed previously in our study of the UGGT:SEP15 complex (8), the structure of human SEP15 U65C predicted by AlphaFold3 (AF3) shows a two-domain protein: Residues 1-45 form a helical bundle of three helices connected by three disulfide linkages, and residues 57-134 adopt a thioredoxin-fold in close agreement with the NMR structure of the homolog from *D. melanogaster* (3) (note that all amino acid numbering reflects the mature form of SEP15 following cleavage of the signal peptide). The modeled structure of the CRD contains three helices connected by relatively short linkers. Interestingly, a mixture of both α and 3_10_ helical configurations is present (Fig. 1A). Helix 1 (h1) is an α-helix that spans residues S7 to E12. Helix 2 (h2) is purely 3_10_ and includes residues C21 to F30. Finally, helix 3 (h3) is a mixture of α (D36 to C42) and 3_10_ (L33 to L35) configurations. Three disulfide bridges connect the helices: C10-C42 connects h1 to h3, while C21-C43 and C24-C39 connect h2 to h3 (Fig. 1B). Residues F5 to Q44 are predicted with C_α_ pLDDT scores above 80 (Fig. S1), indicating confidence in the peptide backbone. The remaining residues are found close to either terminus and are expected to have significant internal mobility.

**Figure 1.**
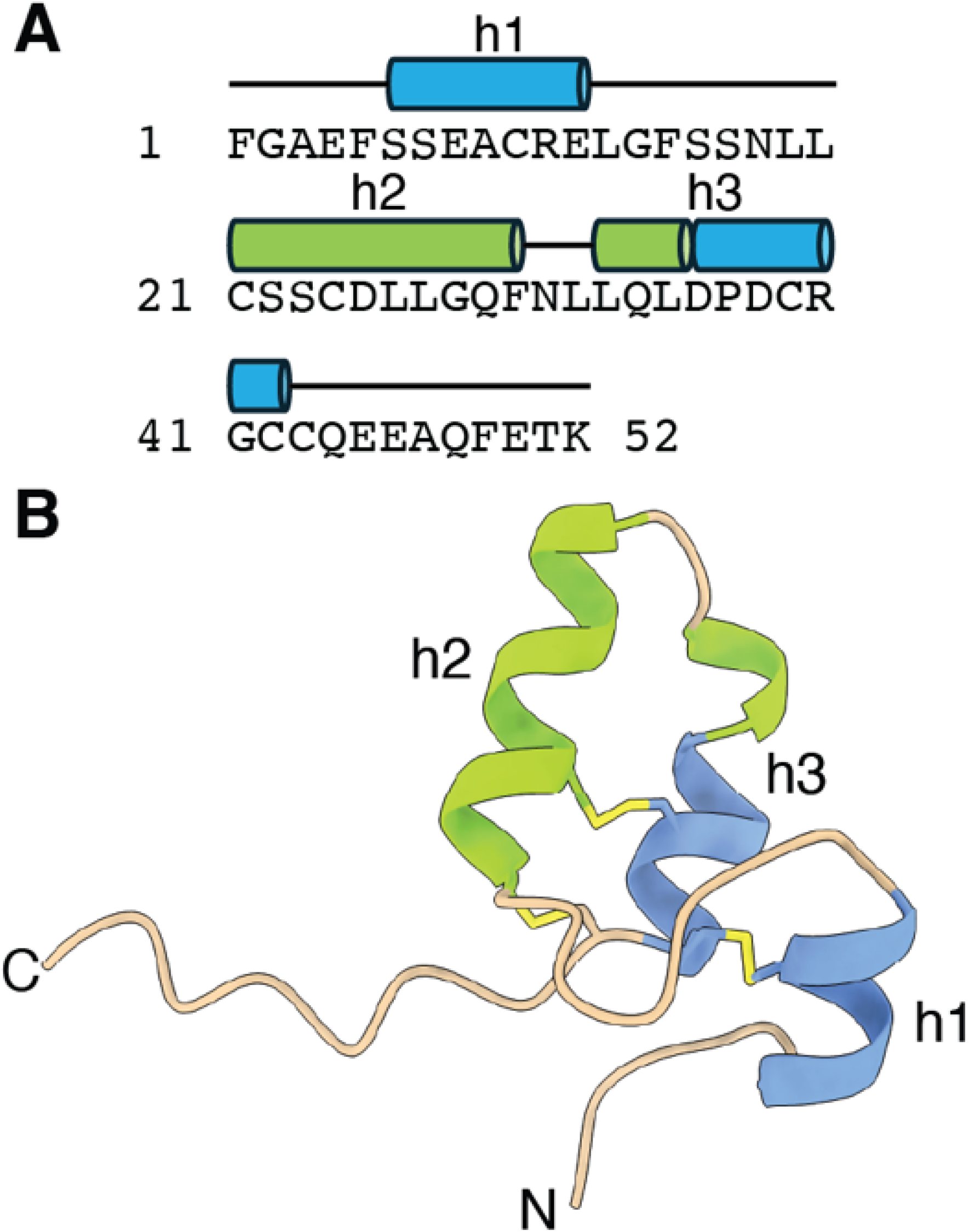
AlphaFold3 prediction of the SEP15 CRD structure. A) The amino acid sequence of the CRD construct is shown annotated with secondary structure assignments. Helices are represented as blue cylinders for α and green for 3_10_. Regions without secondary structure assignment are indicated with a black line. B) Residues 1-52 from the AlphaFold3 prediction are shown as a ribbon diagram colored by secondary structure: blue for α-helix, green for 3_10_-helix, and tan for loop or unstructured regions. Both termini are labelled and side-chain heavy atoms of Cys residues are shown to highlight disulfide bonds.

Strikingly, FoldSeek (10) revealed no structurally homologous domains to the CRD. The only results were CRDs from other SEP15 homologs (Table S1). In contrast, searches using full-length SEP15 as the input found many matches to the thioredoxin domain. Based on this result, we conclude that the CRD adopts a previously undescribed fold.

### Recombinant SEP15-CRD from E. coli SHuffle cells is well folded and retains its UGGT binding function

Using the predicted structure as a guide, we designed a CRD construct including residues 1 to 52. The CRD construct was expressed in *E. coli* SHuffle cells, a bacterial strain that has been engineered to promote the formation of disulfide bonds (11).

To assess whether the *E. coli* expressed CRD was stably folded, we prepared an isotope-labelled sample for NMR spectroscopy. The ^1^H,^15^N-HSQC spectrum of SEP15-CRD (Fig. 2A) reveals 47 well-dispersed backbone amide signals, close to the 50 expected from the sequence. This result shows that CRD from SHuffle cells is folded and adopts a single conformation.

**Figure 2.**
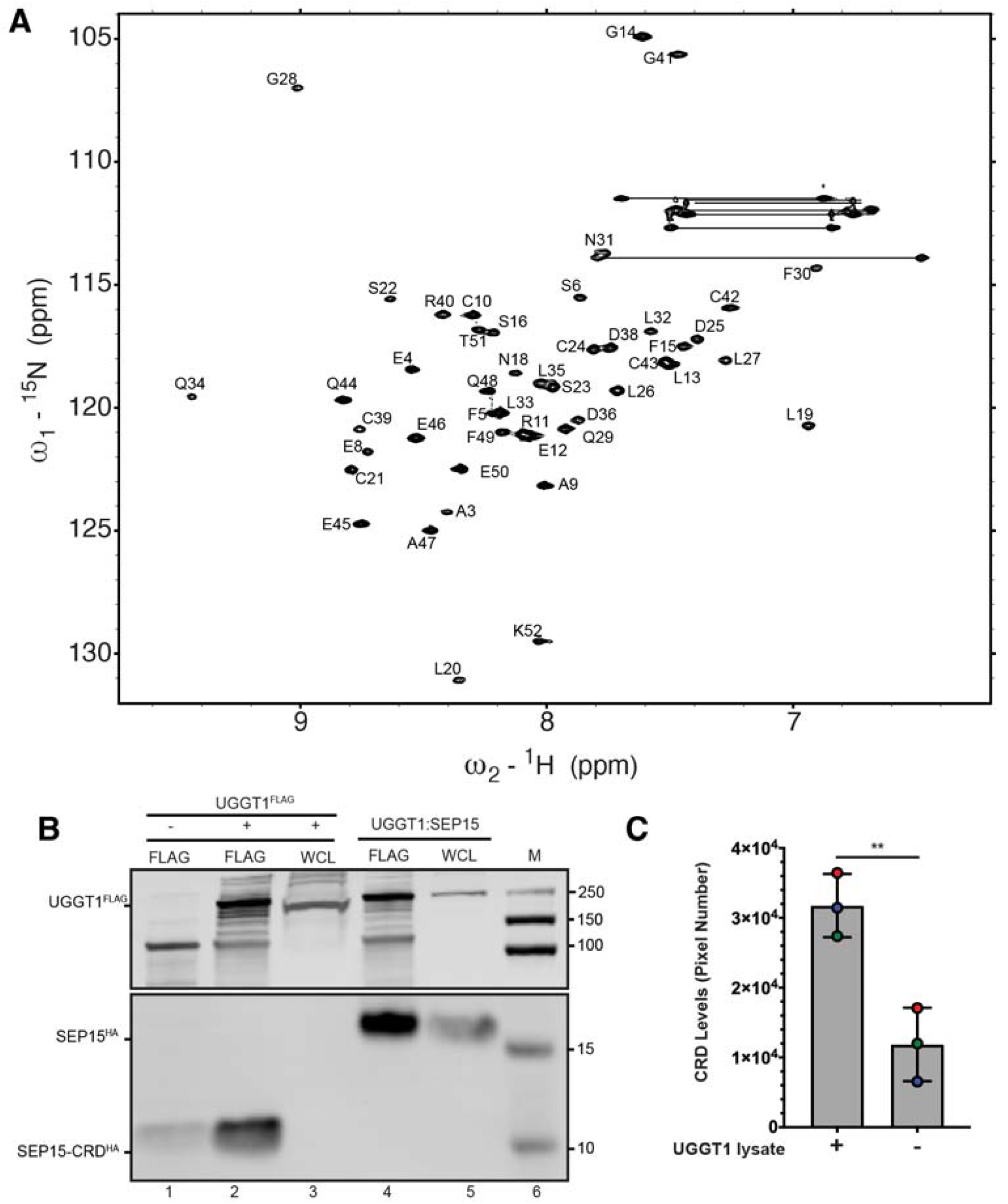
Recombinant CRD construct is folded and functional. A) ^1^H,^15^N-HSQC spectrum of recombinant CRD. The dispersion of backbone amide resonances is characteristic of a folded protein. B) Recombinant CRD from SHuffle cells co-immunoprecipitates (IPs) with UGGT1. Representative western blots are shown with the following lanes: 1 – FLAG pulldown from HEK lysate in the absence of UGGT1^FLAG^ overexpression and addition of bacterial SEP15-CRD^HA^, 2 – FLAG pulldown from HEK lysate with overexpressed UGGT1^FLAG^, 3 – whole cell lysate (WCL) from HEK cells overexpressing UGGT1^FLAG^, 4 – FLAG pulldown from HEK lysate with co-expression of UGGT1^FLAG^ and SEP15^HA^, 5 – WCL from the same sample as lane 4, 6 – molecular weight marker (M). SEP15-CRD^HA^ is enriched in the presence of UGGT1^FLAG^. The same membrane was probed twice, once with aFLAG (upper) and reprobed with aHA (lower) antibodies. Cropped regions of the full image containing the dominant bands are shown. Uncropped images are included in the SI (Figs. S6 and S7). C) Quantification of CRD co-immunoprecipitation. SEP15-CRD^HA^ bands from panel B lanes 1 and 2 were quantified by densitometry. Bars represent the average of three biological replicates (data shown as filled circles) and error bars show the standard deviation. Significantly more SEP15-CRD^HA^ is observed in the presence of UGGT1 lysate (**p < 0.01, two-tailed t-test).

Next, we tested the function of the recombinant CRD by assessing its ability to bind to UGGT1 in a co-immunoprecipitation assay. In a control experiment, UGGT1^FLAG^ and SEP15^HA^ were overexpressed in HEK cells. A FLAG pulldown captured UGGT1 and SEP15^HA^ as well (Fig. 2B, lane 4). Then, recombinant CRD bearing an HA-tag (CRD^HA^) was added to lysate from HEK cells overexpressing UGGT1. Pulling down UGGT1 also led to a significant band for CRD^HA^ (Fig. 2B lane 2 and Fig. 2C), demonstrating that the recombinant domain retained the ability to bind UGGT1.

### The disulfide pattern in the recombinant, functional CRD is consistent with the AlphaFold3 prediction but differs from previous reports

The recombinant CRD was analyzed by mass spectrometry (MS), and a monoisotopic mass of 5732.48 Da was observed, consistent with the expected amino acid sequence and three disulfide bonds. To shed light on the pattern of disulfide bonds, the recombinant CRD was digested with a combination of trypsin and chymotrypsin. Attempts at definitively determining the disulfide linkages by peptide mapping were stymied by the close spacing of Cys residues in the sequence this small domain. MS analysis of the proteolytic digest found four peptides connected by three disulfide bonds: [6-11], [21-30], [34-40], and [41-49] (Table S2). Of the 15 theoretically possible disulfide-bonding patterns for the CRD, only 8 are consistent with the MS data. The pattern predicted by AF3 falls within these 8 options, but the counterpart from the literature does not (9).

### NMR spectroscopy provides support for the AF3-predicted structure of the CRD

Using standard triple-resonance techniques, backbone resonance assignments were determined for SEP15-CRD. The backbone chemical shifts were used to predict backbone torsion angles (Table S3) with the TALOS-N software (12). This tool has a reported accuracy of < 3.5 % and is commonly used to generate backbone torsion restraints for NMR solution structure determination. The results show a striking agreement to the torsions in the AlphaFold prediction (Fig. 3A, Table S4). Comparison of the two sets of ϕ/φ angles in Ramachandran space shows that the same regions are represented. A cluster of residues occurs in the helical region of the diagram near (−60°, −30°). Interestingly, two residues are found in the first quadrant of the Ramachandran map: G14 and N31. Both of these residues are located in loop regions in the predicted structure. Pairwise comparison of all torsion angles shows a strong correlation between the TALOS-derived dihedral angles and those in the AF3 model, with Pearson R of 0.9 and 0.9 for ϕ and φ, respectively (Fig. S2). The RMSD between TALOS-N and AF3 torsion angles is 7.4° for ϕ and 7.7° for φ. These values are similar to the 12° RMSD found in the original test set for TALOS-N compared to X-ray crystal structures (12). These findings show that the secondary structure of CRD is consistent with the AlphaFold prediction.

**Figure 3.**
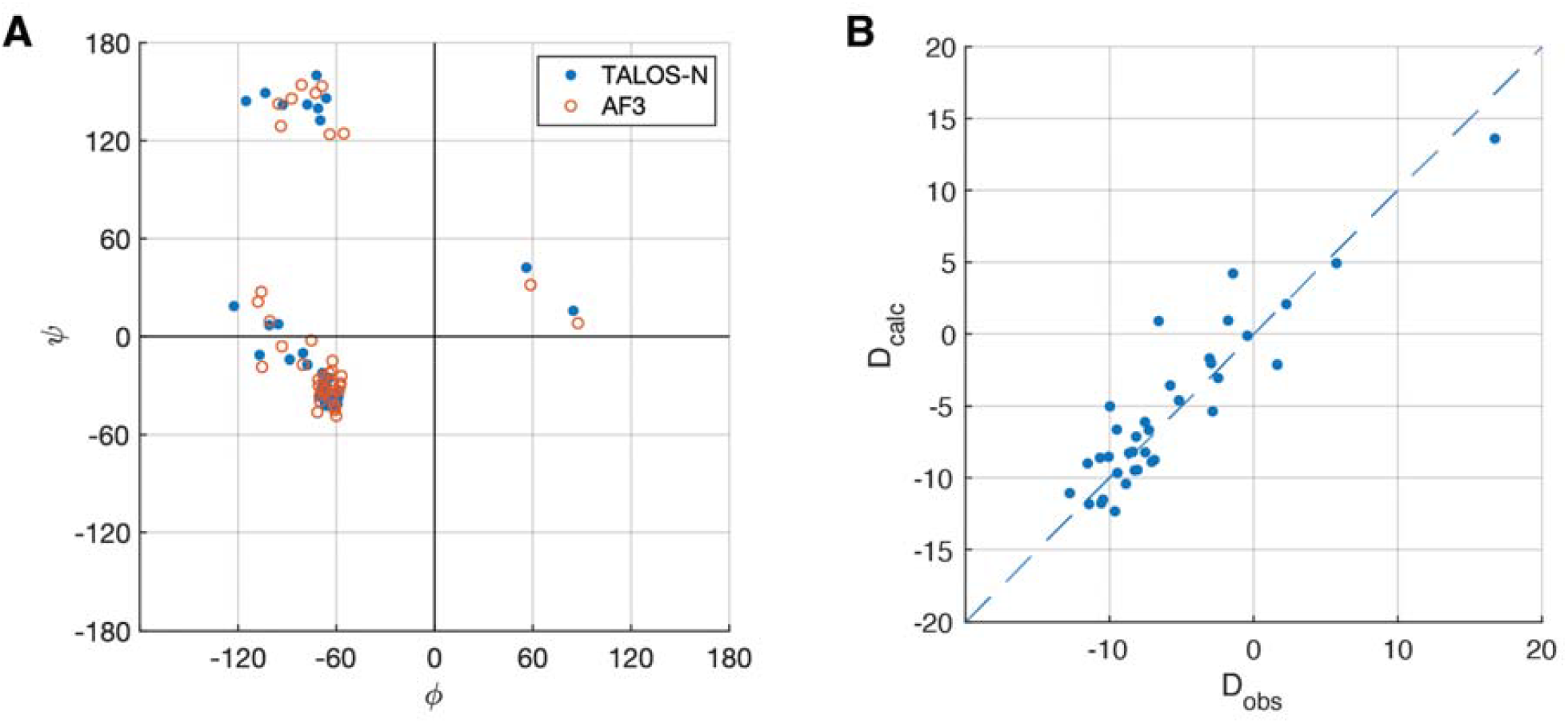
NMR structural restraints agree with the structure predicted by AlphaFold3. A) Ramachandran diagram shows close agreement between backbone torsion angles from TALOS-N (blue, closed circles) and AlphaFold3 (red, open circles). B) Correlation plot shows ^1^H-^15^N RDCs fit well to the AlphaFold3 CRD structure. Experimentally measured couplings (D_obs_) are plotted against back-calculated values (D_calc_).

Hydrogen-deuterium exchange was also monitored by NMR. After 12 minutes of deuterium exchange, a majority of the NH resonance peaks have disappeared from the HSQC spectrum. The remaining protected signals are primarily located within helices h2 and h3 (Fig. S3). These results are consistent with the AF3 model and suggest that this region forms the stable core of this small domain.

To assess the agreement of the global fold relative to the AlphaFold prediction, we measured amide ^1^H-^15^N residual dipolar couplings (RDCs) using phage-oriented media. RDCs are sensitive reporters of amide bond vector orientation and are a convenient method of testing the agreement of candidate structures (13, 14). These values were fit to the AF3 structure and show good agreement, with a Q metric of 0.297 (See Fig. 3B and supporting dataset in SI). Thus, the relative orientation of N-H bond vectors within the CRD is consistent with the AF3 prediction and provides further support that the protein adopts the predicted fold.

### CS-Rosetta discriminates disulfide-bonding pattern

At this stage, the CRD structure emerging from the NMR data is in close agreement with the AlphaFold prediction in both secondary structure and fold, yet the disulfide-bonding pattern was still unverified. We used CS-Rosetta to generate CRD structures for the eight possible disulfide pairings consistent with the MS data (15–18). Only a single disulfide pairing led to an ensemble of structures satisfying the convergence criteria within CS-Rosetta: the pairing predicted by AlphaFold3 (Fig. 4A and table S5). An additional CS-Rosetta calculation was also performed using the literature disulfide pairing. This run did not converge, as shown in Fig. 4B. Based on these results, the AlphaFold-predicted disulfide pairing is the only mode compatible with the observed NMR data.

**Figure 4.**
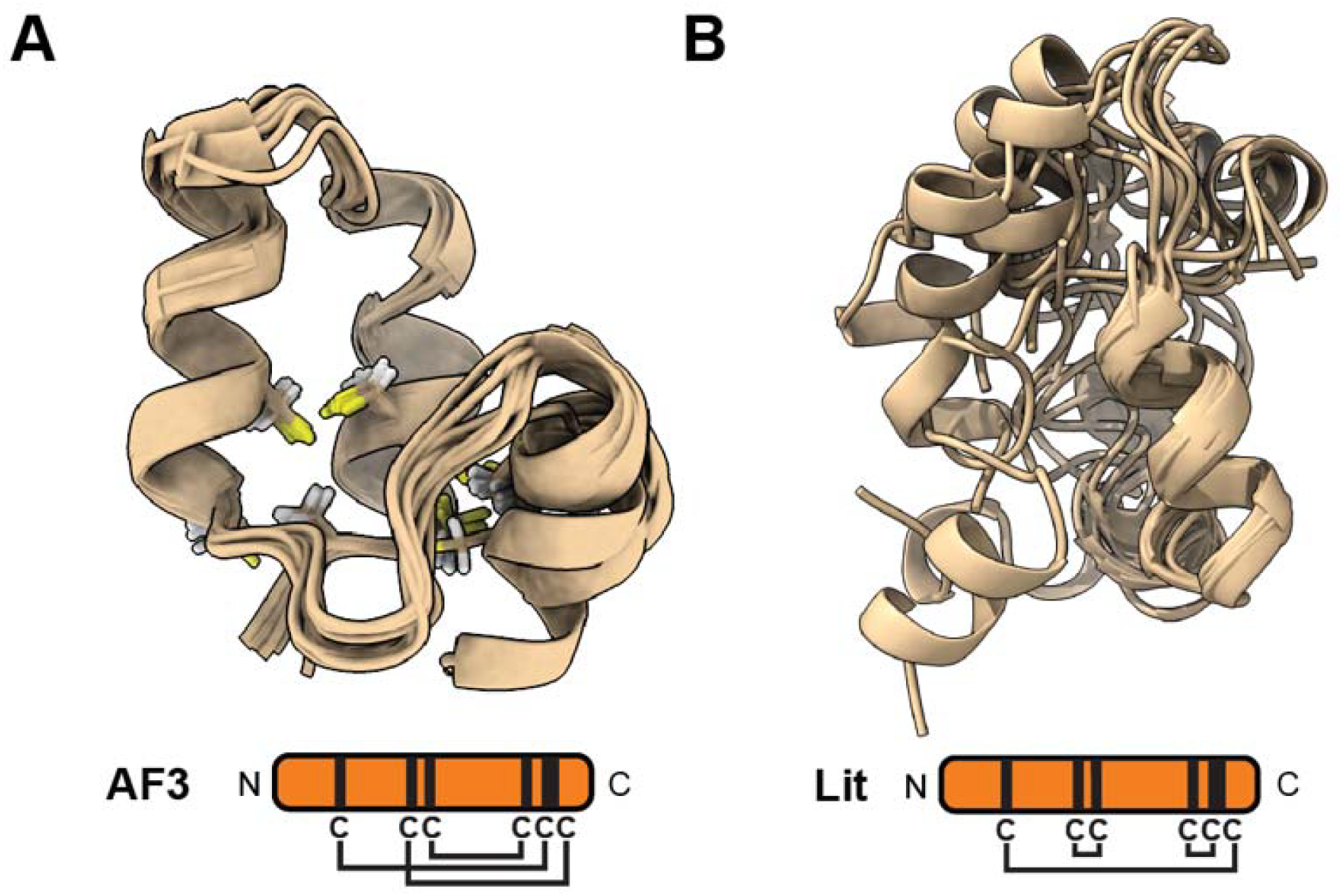
CS-Rosetta only converges for the AlphaFold-predicted disulfide pairing and not alternative pairings. A) CS-Rosetta bundle of top 10 structures determined using the AF3-predicted disulfide pattern: C10-C42, C21-C43, C24-C39. One structure shows an alternate orientation of helix h1. The average C_α_ RMSD within the bundle was 0.9 Å. C) CS-Rosetta result using the literature disulfide pattern (C10-C43, C21-C24, C39-C42) is shown as an example of a non-converged result. The average C_α_ RMSD within the top 10 structures was 2.3 Å.

The CS-Rosetta model closely resembles the AF3 prediction, but the orientation of helix h1 is shifted relative to h2 and h3. Additionally, the top 10 models show variability in this orientation. Repeating the CS-Rosetta calculation with the addition of amide RDC restraints (measured above) led to a tighter ensemble of models and less variability in the h1 position. Nonetheless, the RDC-constrained CS-Rosetta model positions h1 in a different orientation relative to AlphaFold (Fig. S4A). This discrepancy is likely of minor significance. Both models display a suitable hydrophobic surface for interacting with UGGT (Fig. S4B, C). The AlphaFold prediction of the CRD shows little change when predicted as a monomer or in complex with UGGT (Fig. S5B).

Superimposing the CS-Rosetta CRD structure onto CRD in the complex (Fig. S5C) shows the altered position of h1 places the adjacent loop too close to the UGGT SEP15-binding region. It is tempting to speculate that the CS-Rosetta model reveals an alternate conformation of free CRD, but further experiments would be required to assess this in greater detail.

Looking more closely at the secondary structure, CS-Rosetta shows some variability in the helical geometry within the CRD. Helix h1 spans residues S7 to E12 in all ten models and six of the ten also include L13. Next, helix h2 shows more variability. Residues C21 to S23 are assigned as α-helical in 8 of 10 models, while C24 to Q29 are assigned 3_10_ 70% of the time, on average. Finally, helix h3 shows little variation in the CS-Rosetta results. Residues L33 to L35 are assigned as 3_10_-helix in all 10 models, and residues D36 to C39 are α-helical in 9 or more models.

Comparison of the AlphaFold and CS-Rosetta secondary structures show significant agreement. Yet, CS-Rosetta classifies residues C21 to S23 as α-helical while the AF3 prediction places these in a 3_10_-helical configuration. This difference is driven by a change in the hydrogen bonding pattern: CS-Rosetta shows an H-bond between C21O and D25H, whereas AF3 shows an H-bond between C21O and C24H. The HDX results show that C24 amide hydrogen is protected from deuterium uptake, but D25 is more labile. This better fits the AF3 model, where D25 is not participating in a hydrogen bond.

### The structure determined for the isolated CRD is retained in full-length SEP15

Comparison of HSQC NMR spectra of SEP15 constructs encoding the TRX domain and the full-length protein with that of the CRD construct shows that full-length SEP15 overlays well with each single-domain counterpart (Fig. 5). These data show that the full-length protein behaves as a simple combination of the two isolated domains.

**Figure 5.**
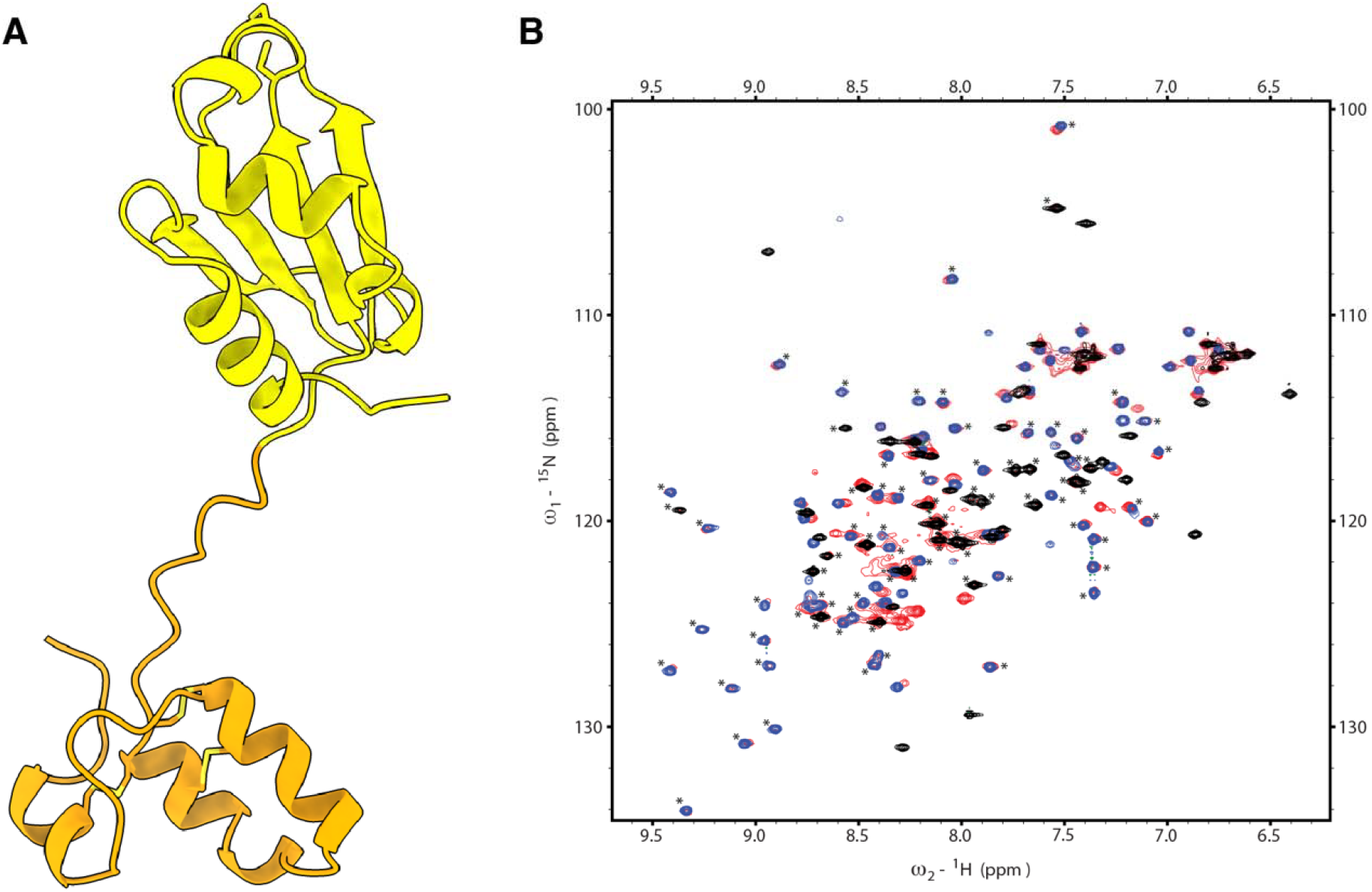
Full-length SEP15 NMR spectrum overlays well with CRD and TRX. A) AF3 prediction of full-length SEP15. The CRD is orange and TRX is yellow. Cys residues are shown in stick representation. B) Overlay of HSQC spectra from CRD (black), TRX (blue), and full-length (red) SEP15 constructs. Backbone amide signals indicated with asterisks overlay well.

## Discussion

We have determined the structure of SEP15 CRD using a combination of AlphaFold prediction and NMR spectroscopy. This small domain is rich in disulfides – three for only ~40 amino acids of well-defined structure. This density is reminiscent of small disulfide-rich toxic peptides found throughout nature (*e*.*g*., conotoxins, defensins, snake venom toxins, *etc*.). Like these small proteins, the CRD structure is likely largely governed by the disulfide bonds as this domain is too small to form a hydrophobic core. It is noteworthy that the CRD contains an unusually long 3_10_-helix of 10 residues. These helices are generally less stable than α-helices because of their less favorable hydrogen bond geometry, and typically short segments on one end of an α-helix adopt the 3_10_-helical fold. We speculate that the long 3_10_-helix within the SEP15 CRD is stabilized by the constraints of disulfide bonds C21-C43 and C24-C39, which anchor it to helix h3. Of course, this speculation raises provocative questions about the folding of the SEP15 CRD and the cooperative formation of its secondary structure and disulfides.

A natural question emerging from our work is why the disulfide pairing mode determined here differs from that previously reported in the literature (9). We suspect that the approach used in the earlier study, which relied on chemical synthesis of SEP15, may have led to an alternatively folded species. Perhaps a kinetic product formed during the oxidative refolding protocol employed in that study. Instead, our protocol leverages the *E. coli* SHuffle strain to promote disulfide bond formation. The evidence we have been able to obtain experimentally (both MS and NMR) and the predicted AF3 structure are not consistent with the pairing reported previously. Most importantly, we have shown that our CRD construct interacts with UGGT1.

SEP15 plays an important role in ER protein quality control through its interaction with UGGT. Our data confirm the predicted structure of CRD and bolster our earlier report predicting the structure of the SEP15/UGGT1 complex (8). This model places the TRX domain and reactive C-G-U motif close to the catalytic site of UGGT1. In this position, SEP15 is poised to reduce or isomerize disulfides close to a N-glycan within UGGT1 substrates. These data lay the groundwork for future studies aimed at understanding the functional interaction of SEP15/UGGT complexes with their substrates.

## Materials and Methods

### Structure prediction

The structure of SEP15 U65C was predicted with the AlphaFold3 server (alphafoldserver.com) from Uniprot accession O60613 using residues 32-165. The single Sec residue was changed to Cys as this amino acid is not currently supported by AlphaFold. The top ranked structure is available at Model Archive under the deposition ma-asb07. All structural analysis and visualization was performed in UCSF ChimeraX (19). Secondary structure assignments were calculated using DSSP as implemented in ChimeraX (20).

### Cloning

A synthetic gene encoding SEP15 U65C (Uniprot O60613, residues 32-165, Azenta) with an N-terminal SUMO tag was inserted into a pET29 vector using Gibson assembly. A SEP15-CRD construct was generated by deleting the region encoding the thioredoxin domain was using Gibson Assembly (New England Biolabs) and the following primers: 5’-CGAGACCAAGTAATAGAATTCGAGCTCCGTCGACAAG-3’ and 5’-CGAATTCTATTACTTGGTCTCGAACTGCGCTTC-3’.

A second construct with a C-terminal HA tag was generated by inserting a synthetic gene encoding SUMO-SEP15-His-HA (IDT, gblock) into a pET29 vector by Gibson assembly using the following oligonucleotide primers: 5’-AATTCGAGCTCCGTCGACAAG-3’ and 5’-ATGTATATCTCCTTCTTAAAGTTAAACAAAATTATTTCTAGAGG-3’.

The resulting vector was then used to generate a truncated CRD-HA construct by deletion of the TRX domain using the following oligonucleotide primers (IDT): 5’-GTGGTGGTGATGGTGCTTGGTCTCGAACTG-3’ and 5’-CAGTTCGAGACCAAGCACCATCACCACCAC-3’.

### Expression and purification of CRD

*E. coli* SHuffle cells (NEB) were transformed with the pET29-SUMO-SEP15-CRD plasmid and positive transformants were selected by kanamycin resistance. One colony was used to inoculate 10 mL of LB media and incubated overnight at 30 °C. The following day, the overnight culture was added to 1 L of LB media and grown at 30 °C until an O.D. 600 reading of 0.8. Expression was induced by addition of IPTG to a final concentration of 0.5 mM. The temperature was lowered to 18 °C and the culture incubated overnight. After 20 hours of expression, cells were harvested by centrifugation and resolubilized in lysis buffer (20 mM Tris, 500 mM NaCl, pH 8.0). Resolubilized cells were flash frozen in liquid nitrogen and stored at −80 °C until further use.

Frozen cells were thawed, and the buffer was supplemented with protease inhibitors (Cocktail Set V, Sigma). Cells were lysed by sonication and centrifuged at 20,000 rpm for 30 minutes (Beckman 25.5 rotor). The resulting supernatant was applied to a 5 mL HisTrapFF column (Cytiva). The column was then washed with buffer A (20 mM Tris, 500 mM NaCl, 20 mM imidazole, pH 8.0). Protein was eluted by a gradient of 0-100 % buffer B (20 mM Tris, 500 mM NaCl, 250 mM imidazole, pH 8.0). Fractions containing the SUMO-CRD fusion protein were pooled and buffer exchanged into buffer A until residual imidazole was less than 20 mM and concentrated to approximately 30 mL (Amicon 10K MWCO, Millipore). The SUMO fusion tag was cleaved by addition of 1 mL of 25 mM Ulp1 protease (see supplementary information) and incubated overnight at 4 °C. CRD was purified from His-SUMO by a second round of Ni-NTA chromatography. Fractions containing the CRD were identified by SDS-PAGE (GenScript) and concentrated to less than 2 mL for size exclusion chromatography. The concentrated solution was applied to a SuperDex 75pg column (Cytiva) equilibrated in buffer C (20 mM sodium phosphate, 100 mM NaCl, pH 7.4).

### Mass spectrometry and peptide digests

CRD and CRD-HA were buffer exchanged into 20 mM ammonium acetate solution using Zeba 7K desalting columns (ThermoFisher). Samples were treated with 375 mM iodoacetamide (Pierce) in ammonium acetate for 30 minutes in the dark. Protein samples were digested overnight with trypsin and chymotrypsin (50:1 protein to protease) at 37 °C.

Samples were diluted 100-fold with 50% aqueous MeOH and 1% formic acid solution prior to mass spectrometry analysis. All mass spectrometry data were collected using a 7 T Bruker SolariX FT-ICR mass spectrometer. Samples were infused at 2 mL/min via syringe pump. Spectra were collected over a range of m/z 150 to 3000, contained 1 million data points, resulting in a 0.7340 s transient, and were the average of 50 scans. Ions were accumulated for 0.3 to 0.5 s prior to trapping in the infinity cell.

The m/z axis was calibrated using 0.2 mg/mL sodium trifluoroacetate clusters in 50 % (v/v) aqueous methanol. Bruker DataAnalysis software was used to deconvolute the spectra, generate peak lists, and simulate isotope distributions. Peptides were assigned using the Protein Prospector MS-Bridge tool (prospector.ucsf.edu) using a mass error tolerance of 5 ppm.

### Cell culture and transfection

HEK293-6E cells were cultured in DMEM (Sigma D5796) containing 10% fetal bovine serum (Gibco 11965118) and cultured at 37 °C, 5% CO_2_ in 10-cm plates. The cells were tested for mycoplasma using a universal detection kit (ATCC, Cat # 30– 012K). For transfection, 3.5E6 cells were added to a 10-cm dish and grown for an additional 24 hrs. The next day the cells were transfected with plasmids encoding for UGGT1 containing a 3xFLAG tag and Sep15 containing an HA tag at a 25:75 ratio, respectively, or UGGT1^FLAG^ alone. 8 μg of total DNA was mixed and diluted in 200 μL Opti-Mem^TM^ (Gibco 31985070). In a separate tube 20 μL of polyethylenimine (PEI, Polyscience 24765) was mixed with 180 μL of Opti-Mem^TM^ and incubated for five minutes at 23° C. The PEI:Opti-Mem^TM^ solution was then added to the diluted DNA mixture and incubated at 20 minutes at 23 °C. All 400 μL of the DNA:PEI solution was added dropwise to each plate before being placed back in the incubator for an additional 24 hours at 37 °C and 5% CO_2_.

### Co-immunoprecipitation assay and Western blot

Twenty-four hours after transfection the cells were washed with 3 mL of PBS. After the wash, the plates were placed on ice, and 1 mL of lysis buffer (20 mM MES, 100 mM NaCl, 30 mM Tris pH 7.5, 0.5% Triton X-100) containing a protease inhibitor cocktail (Thermo 1861278) and 20 mM N-ethylmaleimide (NEM) was added. The cells were scraped off, and the lysate was vortexed for 10 min at 4 °C before centrifuging for 10 min at 20,817xg, 4 °C. For the co- and single UGGT1 transfected plates the clarified lysate was then divided and 20% (200 μL) was precipitated to prepare a whole cell lysate (WCL) fraction by mixing in 1 mL of ice-cold acetone and incubating overnight at −20 °C. For the co-transfected plates 20% of the clarified lysate was mixed with 20 μL of Protein-A-Sepharose® 4B resin (Invitrogen, 101042) and 1μL of αFLAG mouse M2 monoclonal antibody (Sigma F1804) to immunoprecipitate UGGT1^FLAG^:Sep15^HA^ complex.

For the single UGGT1^FLAG^ transfected plates 20% of the lysate was taken and 10 μM of recombinantly produced SEP15-CRD containing an HA tag was spiked into the lysate before 20 μL of Protein-A resin and 1 μL of α3xFLAG antibody was added to immunoprecipitate the complex. The next day the WCL lysate samples were centrifuged at 20,817xg, 4 °C for 10 minutes. The acetone was removed, and the resulting pellets were dried at 23 °C for at least 1 hr. The immunoprecipitation samples were washed 3 times with cold PBS at 500xg, 4 °C. After the final wash, 35 μL of gel loading buffer (30 mM Tris-HCl pH 6.8, 9% SDS, 15% glycerol, 0.05% bromophenol blue) with 100 mM dithiothreitol (Sigma D9779) added was mixed to all samples before being treated for 10 min at 95 °C and vortexed for 10 min. Processed samples were loaded on SurePAGE, Bis-Tris, 12% polyacrylamide gels (GenScript) and run with Tris-MES running buffer (50 mM Tris, 50 mM MES, 0.8 mM EDTA, 0.1 % (w/v) SDS). Protein bands were electrophoretically transferred to immobilon-FL PVDF membrane (Millipore) by application of 90 V for 30 min. Membranes were blotted with mouse anti-FLAG M2 monoclonal antibody and goat anti-mouse IRDye-800CW secondary (Li-Cor). Subsequently, membranes were reprobed using anti-HA antibody (Cell Signaling Technologies, C29F4) and goat anti-rabbit IRDye-680RD (Li-Cor 926-68071).

### NMR spectroscopy

Isotope-labelled CRD was expressed using a dual media approach. A 10 mL LB starter culture was inoculated from a glycerol stock, incubated overnight at 30 °C and expanded to 1 L LB media. Once the culture reached an OD600 of 0.8, cells were pelleted by centrifugation at 3000 rpm (JLA 9.1 rotor) and resuspended in 250 mL of M9 minimal media containing ^15^N-NH_4_Cl and either D-glucose or ^13^C_6_-D-glucose (Cambridge Isotopes) for {U-^15^N} and {U-^15^N,^13^C} samples, respectively. Following media exchange, cells were cooled to 18°C for 30 minutes and expression was induced by addition of IPTG to a final concentration of 0.5 mM. Remaining expression and purification protocols were identical to those described above.

NMR samples were prepared in buffer C supplemented with 5% D_2_O, 10 mM 2,2-dimethyl-2-silapentane-5-sulfonate (DSS) and 0.02 % (w/v) sodium azide. All NMR spectra were acquired using a 600 MHz Bruker magnet equipped with an AVANCE III console and triple-resonance cryoprobe. CRD resonance assignments were determined using a combination of ^1^H,^15^N-HSQC, 3D ^15^N-edited TOCSY-HSQC, HNCA, HN(CO)CA, HNCACB, CBCA(CO)NH, HNCO, and HN(CA)CO experiments. All spectra were processed in NMRpipe and analyzed using NMRFAM-Sparky (21, 22). Proton chemical shifts were referenced directly to the methyl signal of DSS, while ^13^C and ^15^N chemical shifts were referenced indirectly. TALOS-N calculations were performed using the Bax lab webserver (https://spin.niddk.nih.gov/bax-apps/nmrserver/talosn/). CS-Rosetta calculations were performed using the BMRB server (https://csrosetta.bmrb.io).

Residual dipolar couplings were measured on an NMR sample prepared with 15 mg/mL Pf1 bacteriophage using an IPAP-HSQC experiment (23). The measured coupling constants were fit to the AlphaFold structure using PALES software (24). PALES output is included as supporting information. Residues flagged as dynamic by TALOS-N were excluded from the fit and hydrogen atoms were added to the structure using Reduce (25). Correlation plots were generated in MATLAB.

Hydrogen-deuterium exchange experiments were performed by lyophilizing a ^15^N-CRD sample prepared in 20 mM MES, 100 mM NaCl, pH 6, 100% H_2_O buffer and reconstituting with D_2_O. The sample was quickly placed in the spectrometer and ^1^H,^15^N-HSQC spectra were recorded in 10 min intervals.

## Data Availability

The AlphaFold3 prediction of SEP15 is available at model-archive under accession ma-asb07. NMR backbone chemical shifts, resonance assignments, and N-H RDCs have been deposited in the BMRB under accession 52892. Lastly, CS-Rosetta models of SEP15-CRD have been deposited in the PDB-IHM under accession codes 9A9H and 9AHI.

## Acknowledgements

We thank Peter Chien for the generous gift of *E. coli* BL21 (DE3) cells containing the Ulp1 plasmid. Mass spectrometry data were collected using the UMass Amherst Mass Spectrometry Core Facility (RRID:SCR_019063). NMR spectrometry experiments were performed in the bio-NMR core facility at UMass Amherst. This study made use of NMRbox: National Center for Biomolecular NMR Data Processing and Analysis, a Biomedical Technology Research Resource (BTRR), which is supported by NIH grant P41GM111135 (NIGMS). This work was supported by NIH grants R35-GM118161 (LMG), R01-GM086874 (DNH), F32-GM156016 (RVW), and T32-GM008515 (KPG).

