## Supporting Information for "The cysteine-rich domain of SEP15, a selenoprotein co-chaperone of the ER chaperone, UDP-glucose:glycoprotein glucosyltransferase, adopts a novel fold"

University of Massachusetts Amherst

Amherst, MA 01003

**Supporting Information:**

**Supporting methods**

**Supporting Table S1**

**Supporting Table S2**

**Supporting Table S3**

**Supporting Table S4**

**Supporting Table S5**

**Supporting Figure S1**

**Supporting Figure S2**

**Supporting Figure S3**

**Supporting Figure S4**

**Supporting Figure S5**

**Output of PALES software**

***Expression and purification of Ulp1 protease***

A single colony of freshly transformed *E. coli* BL21(DE3) cells containing a pET23-Ulp1 plasmid was used to inoculate 20 mL of culture medium (LB, 100 mg/mL ampicillin), and the culture was incubated overnight at 37 °C. The following day, the overnight culture was expanded to 1 L. Expression was induced at OD600 of 0.6 by addition of IPTG to 0.5 mM concentration and allowed to proceed for 4 hours at 37 °C. Cells were harvested by centrifugation, resolubilized in 50 mM sodium phosphate, 300 mM NaCl, 25 mM imidazole, pH 7.5 buffer (buffer A), and lysed via microfluidizer. Lysate was cleared by centrifugation and the resulting supernatant was applied to a HisTrapFF column (Cytiva). Ulp1 was eluted by a gradient of 0-100% buffer B (50 mM sodium phosphate, 300 mM sodium chloride, 250 mM imidazole, pH 7.5) over 400 mL. Fractions containing Ulp1 were pooled, concentrated, and exchanged into buffer A. Concentrated sample was diluted 1:1 with glycerol and stored at -80 °C until further use.

**Table S1.** FoldSeek matches for SEP15 CRD from AlphaFold Database-Proteome. All entries are direct homologs of SEP15.

| **Target** | **Scientific Name** | **Prob.** | **Seq. Id.** | **E-Value** | **Score** | **Query Pos.** | **Target Pos.** |
| --- | --- | --- | --- | --- | --- | --- | --- |
| AF-A0A077Z1R1-F1-model_v4 | [Trichuris trichiura](https://www.ncbi.nlm.nih.gov/Taxonomy/Browser/wwwtax.cgi?mode=Info&id=36087) | 1 | 55.8 | 1.20E-04 | 202 | 5-47 (52) | 1-43 (125) |
| AF-A0A0N4UDK4-F1-model_v4 | [Dracunculus medinensis](https://www.ncbi.nlm.nih.gov/Taxonomy/Browser/wwwtax.cgi?mode=Info&id=318479) | 1 | 38.2 | 1.06E-03 | 163 | 1-46 (52) | 1-47 (137) |
| AF-A0A0K0E8I3-F1-model_v4 | [Strongyloides stercoralis](https://www.ncbi.nlm.nih.gov/Taxonomy/Browser/wwwtax.cgi?mode=Info&id=6248) | 1 | 40.8 | 1.22E-03 | 157 | 5-52 (52) | 31-79 (159) |
| AF-J9FEU3-F1-model_v4 | [Wuchereria bancrofti](https://www.ncbi.nlm.nih.gov/Taxonomy/Browser/wwwtax.cgi?mode=Info&id=6293) | 1 | 43.4 | 2.00E-03 | 156 | 4-48 (52) | 11-56 (142) |
| AF-A0A5K4FCF6-F1-model_v4 | [Schistosoma mansoni](https://www.ncbi.nlm.nih.gov/Taxonomy/Browser/wwwtax.cgi?mode=Info&id=6183) | 1 | 33.3 | 1.86E-03 | 153 | 4-52 (52) | 19-69 (79) |
| AF-Q9VVJ7-F1-model_v4 | [Drosophila melanogaster](https://www.ncbi.nlm.nih.gov/Taxonomy/Browser/wwwtax.cgi?mode=Info&id=7227) | 1 | 42.5 | 4.66E-03 | 143 | 3-48 (52) | 19-65 (178) |
| AF-Q9N4C6-F1-model_v4 | [Caenorhabditis elegans](https://www.ncbi.nlm.nih.gov/Taxonomy/Browser/wwwtax.cgi?mode=Info&id=6239) | 1 | 30.7 | 3.05E-03 | 142 | 2-52 (52) | 21-72 (152) |
| AF-A0A044SHF7-F1-model_v4 | [Onchocerca volvulus](https://www.ncbi.nlm.nih.gov/Taxonomy/Browser/wwwtax.cgi?mode=Info&id=6282) | 1 | 44.1 | 5.75E-03 | 140 | 7-48 (52) | 34-76 (162) |
| AF-I1JEZ8-F1-model_v4 | [Glycine max](https://www.ncbi.nlm.nih.gov/Taxonomy/Browser/wwwtax.cgi?mode=Info&id=3847) | 1 | 34.6 | 4.66E-03 | 128 | 1-51 (52) | 61-112 (198) |
| AF-C6TB38-F1-model_v4 | [Glycine max](https://www.ncbi.nlm.nih.gov/Taxonomy/Browser/wwwtax.cgi?mode=Info&id=3847) | 1 | 34.6 | 8.78E-03 | 120 | 1-51 (52) | 30-81 (167) |
| AF-C0PKC1-F1-model_v4 | [Zea mays](https://www.ncbi.nlm.nih.gov/Taxonomy/Browser/wwwtax.cgi?mode=Info&id=4577) | 1 | 33.3 | 1.77E-02 | 114 | 2-51 (52) | 26-76 (177) |
| AF-Q8S0K1-F1-model_v4 | [Oryza sativa Japonica Group](https://www.ncbi.nlm.nih.gov/Taxonomy/Browser/wwwtax.cgi?mode=Info&id=39947) | 1 | 44.8 | 2.19E-02 | 110 | 1-47 (52) | 23-71 (162) |
| AF-Q8GWP6-F1-model_v4 | [Arabidopsis thaliana](https://www.ncbi.nlm.nih.gov/Taxonomy/Browser/wwwtax.cgi?mode=Info&id=3702) | 1 | 33.3 | 3.85E-02 | 103 | 1-47 (52) | 26-73 (163) |
| AF-Q4CYI2-F1-model_v4 | [Trypanosoma cruzi strain CL Brener](https://www.ncbi.nlm.nih.gov/Taxonomy/Browser/wwwtax.cgi?mode=Info&id=353153) | 1 | 33.3 | 9.62E-02 | 99 | 4-49 (52) | 51-98 (190) |
| AF-Q387A8-F1-model_v4 | [Trypanosoma brucei brucei TREU927](https://www.ncbi.nlm.nih.gov/Taxonomy/Browser/wwwtax.cgi?mode=Info&id=185431) | 1 | 28.8 | 1.95E-01 | 91 | 1-50 (52) | 76-127 (217) |
| AF-Q4CL23-F1-model_v4 | [Trypanosoma cruzi strain CL Brener](https://www.ncbi.nlm.nih.gov/Taxonomy/Browser/wwwtax.cgi?mode=Info&id=353153) | 0.98 | 37.2 | 3.42E-01 | 79 | 10-50 (52) | 2-44 (135) |

**Table S2.** Observed peptides from CRD digest.

| Peptide | AA seq | Observed Mass | Theoretical Mass | Mass Error (ppm) |
| --- | --- | --- | --- | --- |
| [1-5] | FGAEF | 570.2556 | 570.2563 | -1.3 |
| [16-20] | SSNLL | 533.2930 | 533.2934 | -0.8 |
| [12-15] | ELGF | 465.2340 | 465.2348 | -1.7 |
| [14-20] | GFSSNLL | 737.3837 | 737.3833 | 0.5 |
| [6-11] + [21-30] + [34-40] +  [41-49] |  | 3576.3888 | 3576.3903 | -0.4 |
| [6-11] + [21-30] + [34-49] |  | 3558.3746 | 3558.3798 | -1.5 |
| [6-13] + [21-27] + [34-49] |  | 3468.3565 | 3468.3580 | -0.4 |

**Table S3.** TALOS-N output

| RESID | RESNAME | PHI | PSI | DPHI | DPSI | DIST | S2 | COUNT | CS_COUNT | | CLASS |
| --- | --- | --- | --- | --- | --- | --- | --- | --- | --- | --- | --- |
| 2 | G | 9999 | 9999 | 0 | 0 | 0 | 0 | 0 | 7 | None | |
| 3 | A | 94.416 | -21.871 | 55.464 | 84.606 | 1.522 | 0.413 | 5 | 13 | Dyn | |
| 4 | E | -84.331 | -12.486 | 11.291 | 11.927 | 0.544 | 0.474 | 10 | 17 | Dyn | |
| 5 | F | -91.058 | 141.768 | 15.819 | 10.164 | 0.296 | 0.578 | 8 | 18 | Dyn | |
| 6 | S | -72.147 | 160.008 | 7.962 | 8.072 | 0.145 | 0.704 | 25 | 15 | Strong | |
| 7 | S | -58.8 | -36.852 | 3.552 | 4.067 | 0.108 | 0.822 | 25 | 15 | Strong | |
| 8 | E | -65.573 | -40.495 | 6.596 | 5.103 | 0.088 | 0.872 | 25 | 15 | Strong | |
| 9 | A | -67.074 | -39.038 | 3.217 | 4.831 | 0.074 | 0.885 | 25 | 18 | Strong | |
| 10 | C | -66.002 | -38.133 | 3.035 | 5.079 | 0.075 | 0.884 | 25 | 18 | Strong | |
| 11 | R | -64.228 | -39.515 | 2.69 | 3.23 | 0.063 | 0.879 | 25 | 18 | Strong | |
| 12 | E | -66.441 | -27.018 | 3.022 | 4.369 | 0.071 | 0.839 | 25 | 18 | Strong | |
| 13 | L | -80.394 | -10.144 | 6.183 | 6.536 | 0.088 | 0.793 | 25 | 17 | Strong | |
| 14 | G | 84.69 | 15.809 | 12.26 | 10.672 | 0.136 | 0.738 | 25 | 17 | Strong | |
| 15 | F | -103.409 | 149.293 | 11.565 | 13.406 | 0.215 | 0.733 | 25 | 15 | Strong | |
| 16 | S | -71.168 | 139.801 | 15.389 | 19.239 | 0.251 | 0.717 | 25 | 13 | Strong | |
| 17 | S | -64.271 | -25.174 | 6.897 | 11.067 | 0.289 | 0.737 | 25 | 12 | Strong | |
| 18 | N | -95.495 | 7.605 | 6.401 | 6.121 | 0.268 | 0.761 | 25 | 14 | Strong | |
| 19 | L | -69.848 | 132.49 | 9.632 | 6.921 | 0.253 | 0.819 | 25 | 16 | Strong | |
| 20 | L | -92.801 | 142.016 | 19.757 | 16.814 | 0.331 | 0.866 | 25 | 17 | Strong | |
| 21 | C | -59.77 | -34.806 | 4.19 | 4.823 | 0.403 | 0.88 | 25 | 17 | Strong | |
| 22 | S | -67.119 | -31.237 | 4.148 | 5.684 | 0.371 | 0.886 | 25 | 18 | Strong | |
| 23 | S | -68.368 | -35.318 | 3.892 | 7.236 | 0.455 | 0.886 | 25 | 18 | Strong | |
| 24 | C | -68.669 | -31.687 | 4.667 | 5.033 | 0.511 | 0.888 | 25 | 18 | Strong | |
| 25 | D | -68.62 | -35.143 | 4.611 | 4.861 | 0.409 | 0.883 | 25 | 18 | Strong | |
| 26 | L | -70.768 | -36.743 | 6.843 | 8.401 | 0.347 | 0.864 | 25 | 17 | Strong | |
| 27 | L | -66.477 | -37.623 | 5.816 | 7.364 | 0.357 | 0.857 | 25 | 16 | Strong | |
| 28 | G | -63.811 | -37.117 | 5.365 | 7.186 | 0.243 | 0.849 | 25 | 16 | Strong | |
| 29 | Q | -68.261 | -28.951 | 9.81 | 11.386 | 0.271 | 0.853 | 25 | 17 | Strong | |
| 30 | F | -101 | 6.949 | 9.585 | 7.768 | 0.337 | 0.863 | 25 | 18 | Strong | |
| 31 | N | 56.008 | 42.243 | 5.356 | 8.514 | 0.271 | 0.871 | 25 | 17 | Strong | |
| 32 | L | -122.56 | 18.618 | 9.549 | 11.261 | 0.255 | 0.876 | 25 | 16 | Strong | |
| 33 | L | -61.849 | -38.134 | 6.542 | 9.053 | 0.424 | 0.855 | 25 | 16 | Strong | |
| 34 | Q | -68.667 | -22.166 | 6.871 | 10.302 | 0.25 | 0.864 | 25 | 17 | Strong | |
| 35 | L | -88.543 | -14.021 | 17.455 | 17.448 | 0.269 | 0.882 | 25 | 15 | Strong | |
| 36 | D | -60.846 | -42.903 | 5.429 | 3.843 | 0.24 | 0.922 | 25 | 12 | Strong | |
| 37 | P | -59.332 | -41.175 | 3.985 | 6.044 | 0.22 | 0.929 | 25 | 12 | Strong | |
| 38 | D | -67.02 | -41.173 | 4.756 | 5.488 | 0.28 | 0.926 | 25 | 15 | Strong | |
| 39 | C | -65.912 | -42.277 | 5.023 | 4.295 | 0.308 | 0.914 | 25 | 18 | Strong | |
| 40 | R | -66.291 | -29.221 | 4.496 | 8.448 | 0.302 | 0.894 | 25 | 17 | Strong | |
| 41 | G | -77.77 | -17.176 | 9.349 | 9.259 | 0.396 | 0.888 | 8 | 17 | Warn | |
| 42 | C | -106.943 | -11.317 | 13.365 | 13.315 | 0.381 | 0.888 | 8 | 17 | Warn | |
| 43 | C | -77.762 | 142.151 | 13.952 | 10.581 | 0.421 | 0.887 | 25 | 18 | Strong | |
| 44 | Q | -115.18 | 144.305 | 14.267 | 11.688 | 0.404 | 0.817 | 25 | 18 | Strong | |
| 45 | E | -66.27 | 146.02 | 7.587 | 7.78 | 0.397 | 0.653 | 25 | 18 | Strong | |
| 46 | E | -74.646 | 142.316 | 25.777 | 14.918 | 0.558 | 0.433 | 8 | 18 | Dyn | |
| 47 | A | -74.659 | -20.137 | 12.278 | 14.005 | 0.637 | 0.307 | 4 | 18 | Dyn | |
| 48 | Q | -93.352 | -5.763 | 11.618 | 9.342 | 0.683 | 0.241 | 6 | 17 | Dyn | |
| 49 | F | -79.729 | 140.107 | 10.309 | 14.514 | 1.034 | 0.188 | 25 | 17 | Dyn | |
| 50 | E | -87.704 | 127.863 | 16.407 | 30.62 | 1.042 | 0.113 | 5 | 17 | Dyn | |
| 51 | T | -101.779 | 135.558 | 13.782 | 12.56 | 1.602 | 0.049 | 25 | 17 | Dyn | |
| 52 | K | 9999 | 9999 | 0 | 0 | 0 | 0 | 0 | 11 | None | |

**Table S4.** Backbone torsion angles from AlphaFold3 predicted CRD structure. Secondary structure is indicated by shading: blue for α-helix and green for 3_10_-helix.

| ResNum | ResType | Phi | Psi |
| --- | --- | --- | --- |
| 1 | F | None | 95.4 |
| 2 | G | -149.6 | 23.6 |
| 3 | A | -88.1 | 121.9 |
| 4 | E | -67.5 | -25.1 |
| 5 | F | -101.4 | 147.1 |
| 6 | S | -68.8 | 153.4 |
| 7 | S | -57.6 | -28.4 |
| 8 | E | -71.6 | -46.2 |
| 9 | A | -61.5 | -41.6 |
| 10 | C | -62.9 | -36.0 |
| 11 | R | -70.0 | -35.8 |
| 12 | E | -66.9 | -33.9 |
| 13 | L | -75.3 | -2.4 |
| 14 | G | 87.5 | 8.2 |
| 15 | F | -87.4 | 145.8 |
| 16 | S | -81.3 | 154.2 |
| 17 | S | -62.4 | -14.8 |
| 18 | N | -107.9 | 21.3 |
| 19 | L | -64.1 | 123.9 |
| 20 | L | -95.4 | 142.6 |
| 21 | C | -62.2 | -28.8 |
| 22 | S | -57.9 | -30.1 |
| 23 | S | -70.9 | -29.9 |
| 24 | C | -68.9 | -32.3 |
| 25 | D | -64.5 | -22.0 |
| 26 | L | -80.5 | -17.1 |
| 27 | L | -58.5 | -33.1 |
| 28 | G | -61.0 | -32.8 |
| 29 | Q | -71.0 | -26.3 |
| 30 | F | -100.7 | 9.5 |
| 31 | N | 58.6 | 31.7 |
| 32 | L | -105.9 | 27.4 |
| 33 | L | -56.9 | -24.0 |
| 34 | Q | -62.6 | -20.8 |
| 35 | L | -93.3 | -6.0 |
| 36 | D | -60.7 | -43.2 |
| 37 | P | -60.9 | -44.9 |
| 38 | D | -70.0 | -39.5 |
| 39 | C | -60.1 | -48.5 |
| 40 | R | -66.8 | -32.1 |
| 41 | G | -67.8 | -24.2 |
| 42 | C | -105.4 | -18.5 |
| 43 | C | -72.7 | 149.2 |
| 44 | Q | -93.9 | 128.9 |
| 45 | E | -55.5 | 124.4 |
| 46 | E | -54.5 | 114.6 |
| 47 | A | -47.0 | 111.0 |
| 48 | Q | -42.4 | 108.5 |
| 49 | F | -52.1 | 105.5 |
| 50 | E | -44.1 | 106.0 |
| 51 | T | -49.5 | 105.1 |
| 52 | K | -45.2 | 105.5 |

**Table S5.** Results of CS-Rosetta structure calculations for different CRD disulfide pairing candidates.

| Name | Disulfide Pairing | Converged? | C_a_ RMSD (Å) |
| --- | --- | --- | --- |
| AF3 | C10-C42, C21-C43, C24-C39 | Y | 0.9 |
| 1 | C10-C21, C24-C42, C39-C43 | N | 2.3 |
| 2 | C10-C21, C24-C43, C39-C42 | N | 3.2 |
| 3 | C10-C24, C21-C42, C39-C43 | N | 2.3 |
| 4 | C10-C24, C21-C43, C39-C42 | N | 2.9 |
| 5 | C10-C43, C21-C39, C24-C42 | N | 3.4 |
| 6 | C10-C42, C21-C39, C24-C43 | N | 2.5 |
| 7 | C10-C43, C24-C39, C21-C42 | N | 2.6 |
| Lit | C10-C43, C21-C24, C39-C42 | N | 2.3 |

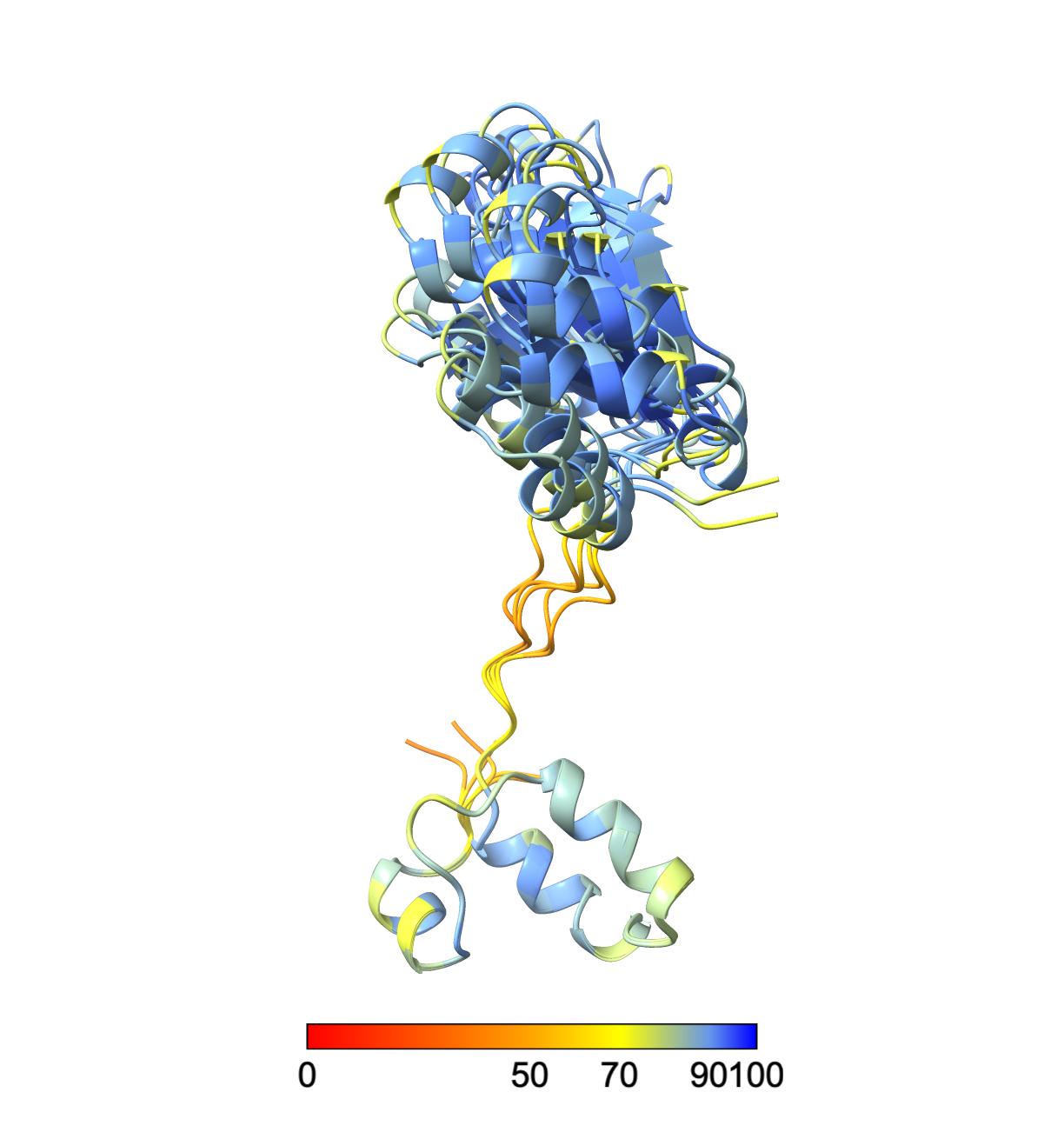

**Figure S1.** AlphaFold prediction of SEP15. All five predicted models are shown colored by pLDDT values. Models aligned by the CRD. The relative orientation between the CRD and TRX domains varies among the models but the structure of each domain is consistent across all models with pairwise backbone RMSDs less than 1 Å.

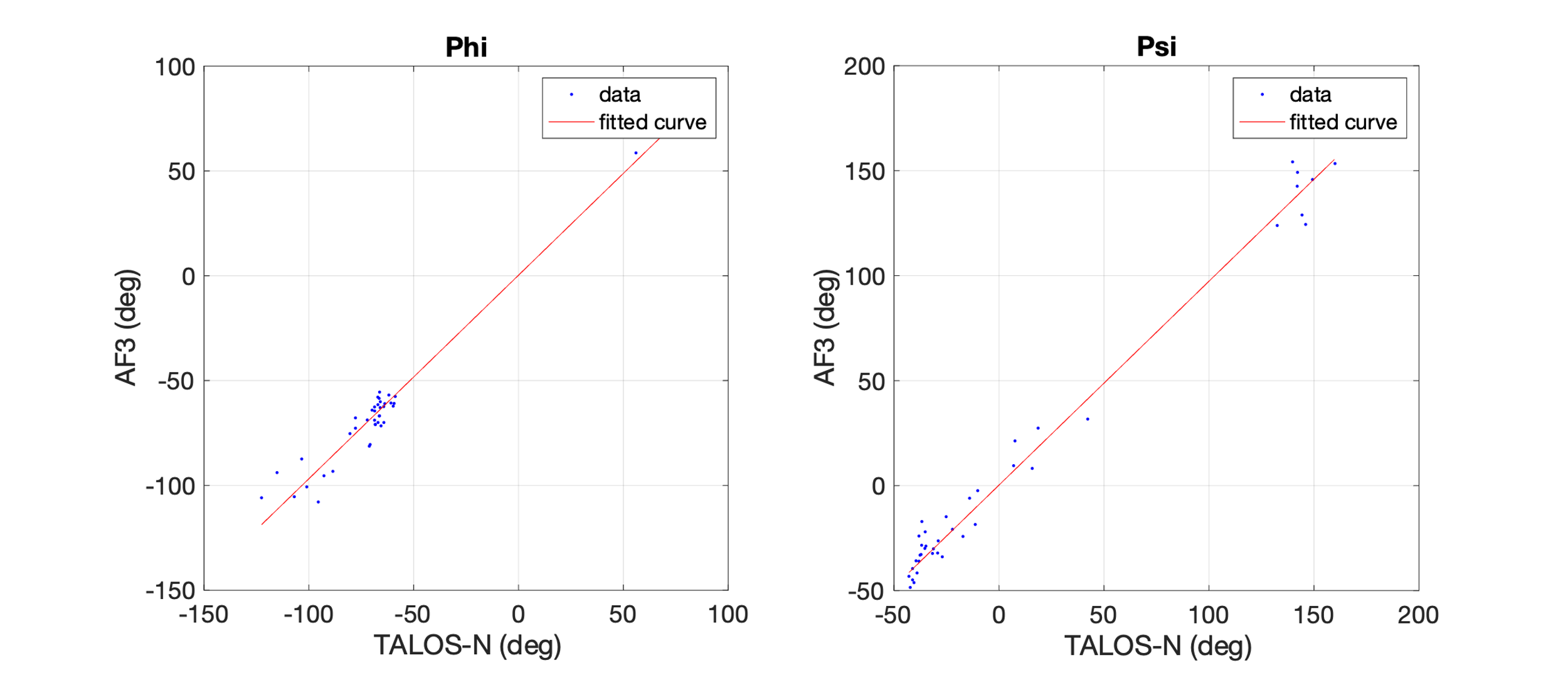

**Figure S2.** Correlation plots comparing backbone torsion angles from TALOS-N and AF3. Phi angles (left) have a line of best fit of y = 0.9710x + 0.2655 and an R^2^ coefficient of 0.9621, while psi angles (right) had a best fit of y = 0.9587x + 1.509 and an R^2^ coefficient of 0.9858. RMSD for Phi and Psi angles was 7.0° and 8.3°, respectively.

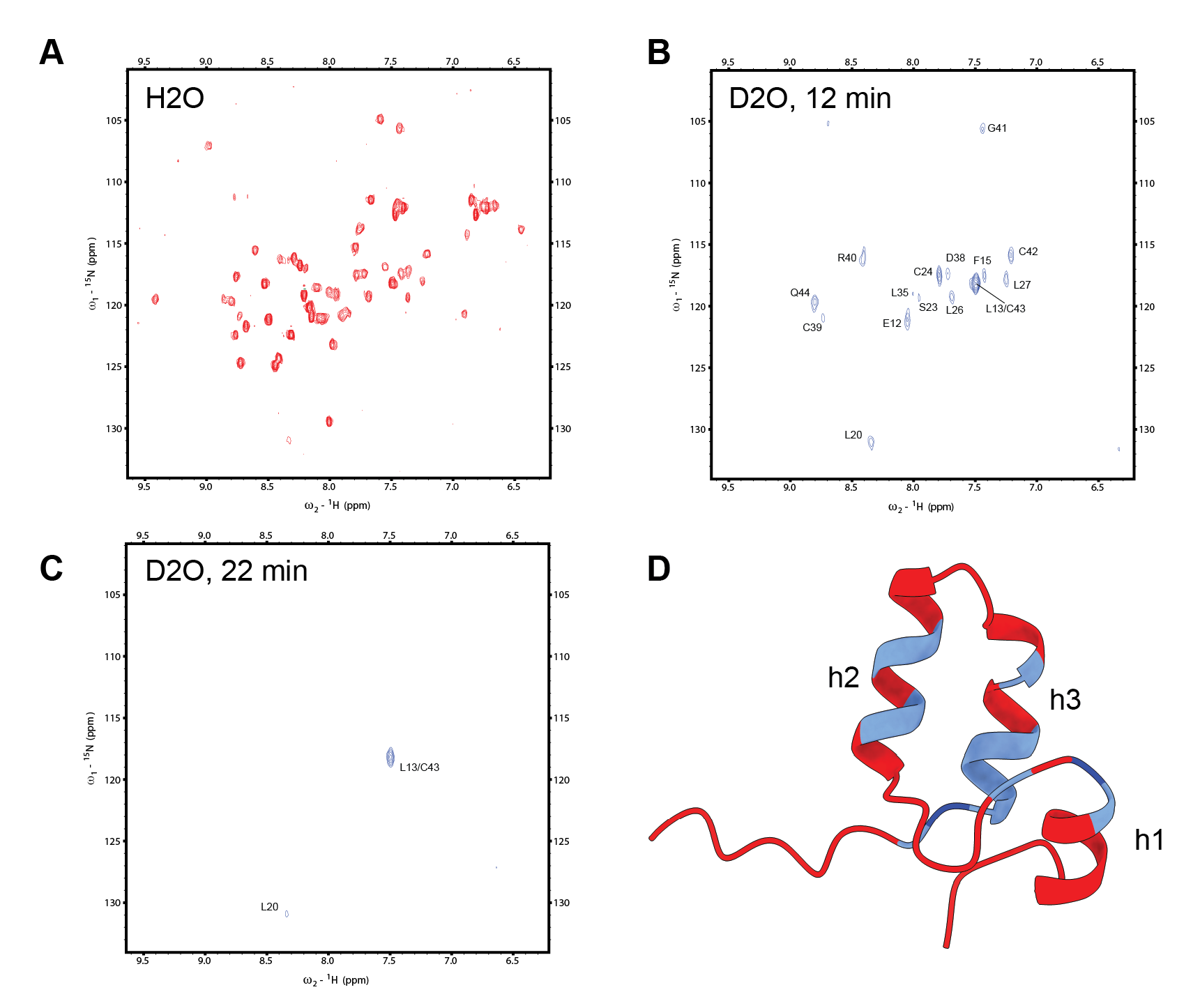

**Figure S3.** Hydrogen-deuterium exchange of SEP15-CRD shows protection in helices h2 and h3. A) 2D NMR HSQC spectrum for CRD acquired in 95/5 % (v/v) H_2_O/D_2_O solution. B) HSQC spectrum acquired after 12 min of deuterium exchange. C) HSQC spectrum acquired after 22 min of deuterium exchange. D) AF3 prediction for SEP15-CRD colored according to HDX data. Red residues exchange too quickly to be observed following transfer into D_2_O solution. Light blue residues exchange at an intermediate rate and were only observed after12 min of deuterium exchange, but not in subsequent spectra. Dark blue residues exchange most slowly and were observed after 22 min of deuterium exchange.

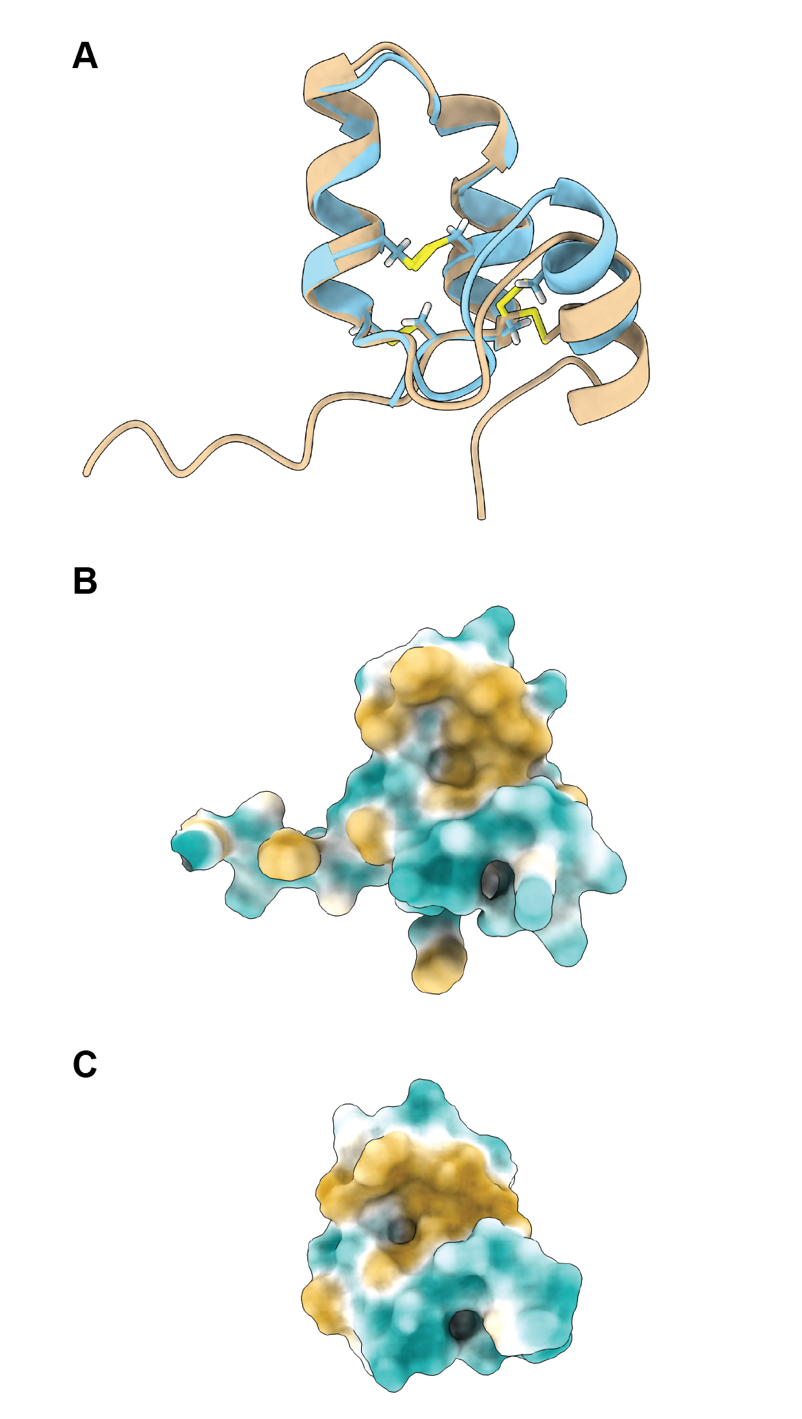

**Figure S4.** AlphaFold3 and CS-Rosetta models of SEP15 CRD are highly similar. A structural overlay (A) of the AF3 (tan) and CS-Rosetta (light blue) models shows that helix h1 has moved relative to the other helices. Hydrophobic surfaces of each CRD model show a solvent-exposed hydrophobic surface patch (gold) in both cases (B and C). These patches are suitable surfaces for interacting with UGGT.

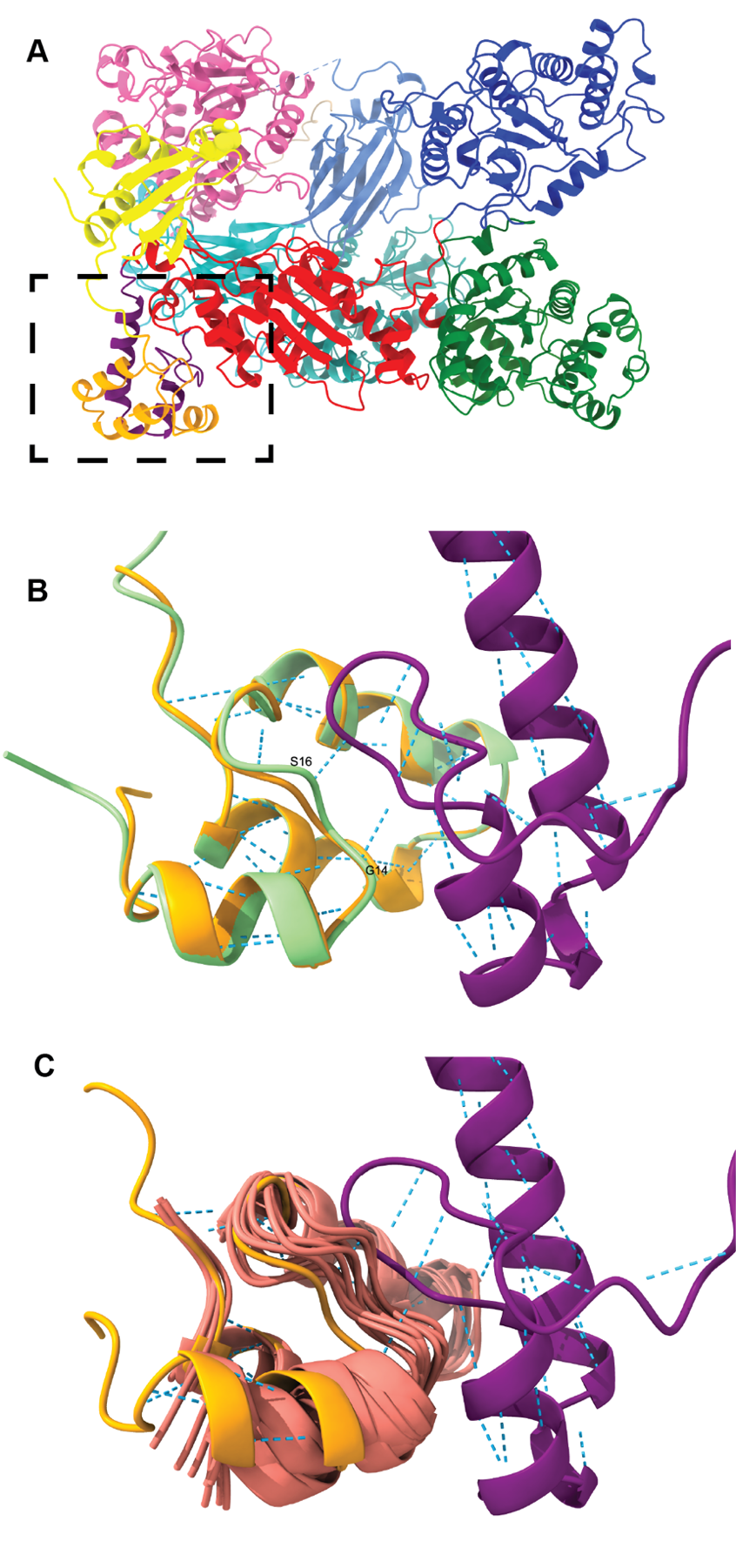

**Figure S5.** The AlphaFold prediction of the UGGT:SEP15 complex shows the SEP15 CRD (orange) in contact with UGGT SEP15-binding region (purple) (A, dashed box). The CRD structure predicted by AF3 (tan) closely matches the CRD within the complex (B). The CS-Rosetta CRD structure (pink) overlays less well, with the largest difference observed in helix h1 (C). Two intermolecular hydrogen bonds are predicted in this region, involving CRD residues G14 and S16.

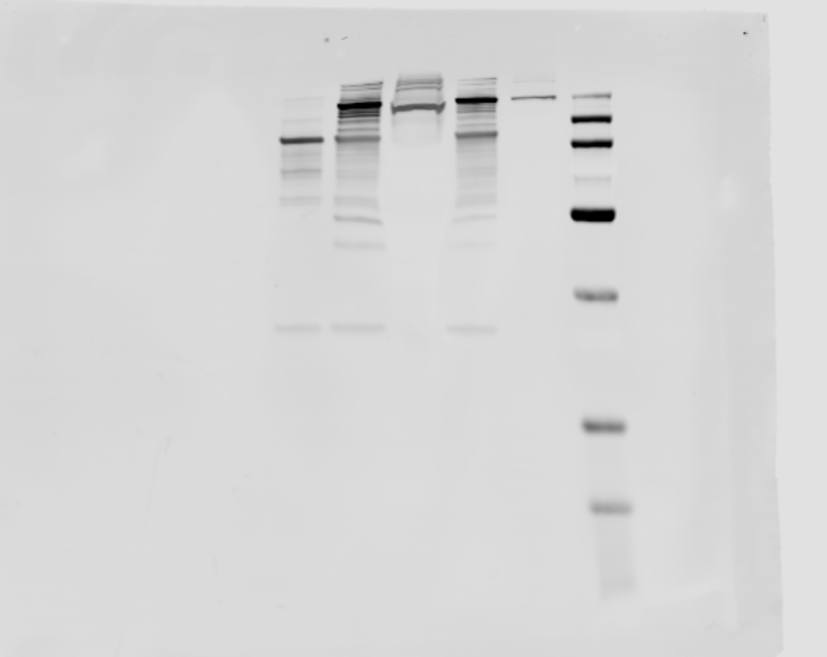

**Figure S6.** Unedited image showing αFLAG Western blot from Fig. 2B.

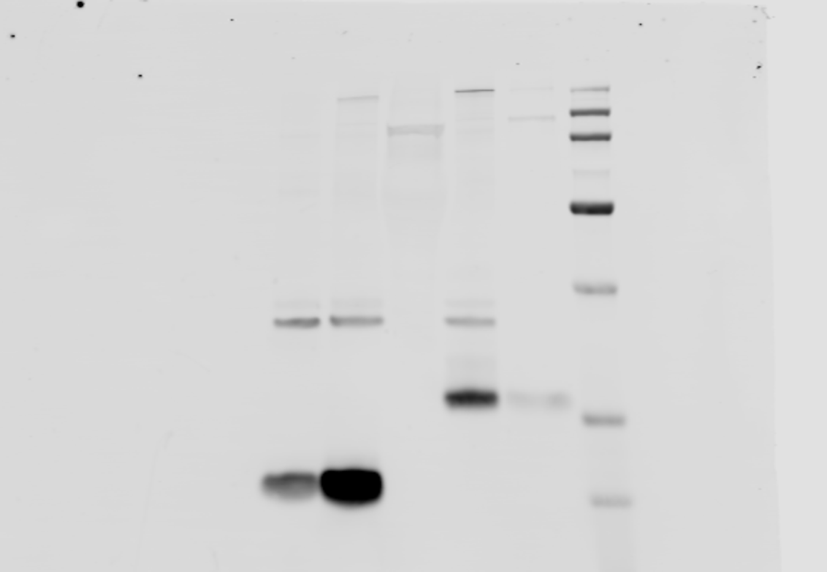

**Figure S7.** Unedited image showing αHA Western blot from Fig. 2B.

**Output from PALES SVD fit using AlphaFold3 prediction**.

```
REMARK Molecular Alignment Simulation.

REMARK Simulation parameters.

DATA PALES_MODE DC

DATA TENSOR_MODE SVD (Order Matrix Method)


REMARK Order matrix.

DATA SAUPE  -1.1140e-04 -8.9729e-05  3.4443e-04  1.4207e-04  1.5798e-05

DATA IRREDUCIBLE REPRESENTATION (A0,A1R,A1I,A2R,A2I)   -3.8010e+00 -3.9581e+00  4.4012e-01 -1.2499e+00 -9.5956e+00
DATA IRREDUCIBLE GENERAL_MAGNITUDE   1.5279e+01


REMARK Eigensystem & Euler angles for clockwise rotation about z, y', z''.

DATA EIGENVALUES (Axx,Ayy,Azz)   -8.8432e-05 -3.3594e-04  4.2437e-04
DATA EIGENVECTORS
DATA EIGENVECTORS XAXIS  1.9016e-01 -4.2070e-01  8.8704e-01
DATA EIGENVECTORS YAXIS  7.2077e-01 -5.5364e-01 -4.1710e-01
DATA EIGENVECTORS ZAXIS  6.6657e-01  7.1868e-01  1.9795e-01

DATA Q_EULER_SOLUTIONS    ALPHA     BETA    GAMMA
DATA Q_EULER_ANGLES  1   334.82    78.58   132.85
DATA Q_EULER_ANGLES  2   154.82    78.58   132.85
DATA Q_EULER_ANGLES  3   205.18   101.42   312.85
DATA Q_EULER_ANGLES  4    25.18   101.42   312.85


REMARK Euler angles (psi/theta/phi) for rotation about x, y, z.

DATA EULER_SOLUTIONS 2
DATA EULER_ANGLES   74.60  -41.80  255.22
DATA EULER_ANGLES  254.60  221.80   75.22

DATA Da  2.121856e-04
DATA Dr  8.250246e-05


REMARK Dipolar couplings.

DATA N                    36
DATA RMS                  2.417
DATA Chi2                 210.257
DATA CORR R               0.912
DATA CORNILESCU Q         0.297
DATA REGRESSION OFFSET    -0.177 +/- 0.563 [Hz]
DATA REGRESSION SLOPE     0.896 +/- 0.069 [Hz]
DATA REGRESSION BAX SLOPE 0.987 +/- 0.053 [Hz]

VARS  RESID_I RESNAME_I ATOMNAME_I RESID_J RESNAME_J ATOMNAME_J DI D_OBS D D_DIFF DD W
FORMAT %4d %4s %4s %4d %4s %4s %9.2f %9.3f %9.3f %9.3f %.2f %.2f

    5  PHE    N    5  PHE    H -38304.92   -7.0700   -8.8902    1.8202  1.0000 1.00
    6  SER    N    6  SER    H -38256.43   -2.9500   -2.0124   -0.9376  1.0000 1.00
    8  GLU    N    8  GLU    H -38279.55  -12.7600  -11.0663   -1.6937  1.0000 1.00
    9  ALA    N    9  ALA    H -38355.26   -7.5200   -8.2197    0.6997  1.0000 1.00
   10  CYS    N   10  CYS    H -38278.86  -10.0600   -8.5337   -1.5263  1.0000 1.00
   11  ARG    N   11  ARG    H -38290.80  -11.4200  -11.8281    0.4081  1.0000 1.00
   12  GLU    N   12  GLU    H -38260.77   -9.4500   -9.6538    0.2038  1.0000 1.00
   13  LEU    N   13  LEU    H -38339.50   -8.3900   -8.1928   -0.1972  1.0000 1.00
   14  GLY    N   14  GLY    H -38245.19   -5.7900   -3.5597   -2.2303  1.0000 1.00
   15  PHE    N   15  PHE    H -38294.07   -7.5400   -6.1047   -1.4353  1.0000 1.00
   18  ASN    N   18  ASN    H -38343.61   -2.8400   -5.3639    2.5239  1.0000 1.00
   19  LEU    N   19  LEU    H -38310.52   -8.1500   -7.1283   -1.0217  1.0000 1.00
   20  LEU    N   20  LEU    H -38234.52   -8.0600   -9.4497    1.3897  1.0000 1.00
   21  CYS    N   21  CYS    H -38340.61   -8.6400   -8.2736   -0.3664  1.0000 1.00
   22  SER    N   22  SER    H -38307.03    2.2700    2.0802    0.1898  1.0000 1.00
   23  SER    N   23  SER    H -38268.17   -5.1800   -4.6039   -0.5761  1.0000 1.00
   24  CYS    N   24  CYS    H -38248.73   -8.2500   -9.4810    1.2310  1.0000 1.00
   25  ASP    N   25  ASP    H -38248.29   -0.4300   -0.1186   -0.3114  1.0000 1.00
   26  LEU    N   26  LEU    H -38270.43   -1.4300    4.2122   -5.6422  1.0000 1.00
   27  LEU    N   27  LEU    H -38303.50  -10.5700  -11.7737    1.2037  1.0000 1.00
   28  GLY    N   28  GLY    H -38276.20   -7.2700   -6.6665   -0.6035  1.0000 1.00
   29  GLN    N   29  GLN    H -38283.85   -3.0600   -1.7015   -1.3585  1.0000 1.00
   30  PHE    N   30  PHE    H -38345.35   -9.6200  -12.3332    2.7132  1.0000 1.00
   31  ASN    N   31  ASN    H -38244.86   -1.7700    0.9369   -2.7069  1.0000 1.00
   32  LEU    N   32  LEU    H -38303.88   -2.4600   -3.0401    0.5801  1.0000 1.00
   33  LEU    N   33  LEU    H -38289.66   -9.4900   -6.6341   -2.8559  1.0000 1.00
   34  GLN    N   34  GLN    H -38272.07   -8.8600  -10.4243    1.5643  1.0000 1.00
   35  LEU    N   35  LEU    H -38355.00  -10.4500  -11.5202    1.0702  1.0000 1.00
   38  ASP    N   38  ASP    H -38272.72  -10.6600   -8.6044   -2.0556  1.0000 1.00
   39  CYS    N   39  CYS    H -38257.78    1.6400   -2.1163    3.7563  1.0000 1.00
   40  ARG    N   40  ARG    H -38350.71    5.7500    4.9263    0.8237  1.0000 1.00
   41  GLY    N   41  GLY    H -38232.55   -9.9700   -5.0155   -4.9545  1.0000 1.00
   42  CYS    N   42  CYS    H -38263.63   -6.8800   -8.7547    1.8747  1.0000 1.00
   43  CYS    N   43  CYS    H -38262.44   16.7400   13.6042    3.1358  1.0000 1.00
   44  GLN    N   44  GLN    H -38256.21   -6.5900    0.9023   -7.4923  1.0000 1.00
   45  GLU    N   45  GLU    H -38312.67  -11.5100   -8.9949   -2.5151  1.0000 1.00
```
